# Mitochondrial PE potentiates respiratory enzymes to amplify skeletal muscle aerobic capacity

**DOI:** 10.1101/599985

**Authors:** Timothy D. Heden, Jordan M. Johnson, Patrick J. Ferrara, Hiroaki Eshima, Anthony R. P. Verkerke, Edward J. Wentzler, Piyarat Siripoksup, Tara M. Narowski, Chanel B. Coleman, Chien-Te Lin, Terence E. Ryan, Paul T. Reidy, Lisandra E. de Castro Brás, Courtney M. Karner, Charles F. Burant, J. Alan Maschek, James E. Cox, Douglas G. Mashek, Gabrielle Kardon, Sihem Boudina, Tonya N. Zeczycki, Jared Rutter, Saame R. Shaikh, Jean E. Vance, Micah J. Drummond, P. Darrell Neufer, Katsuhiko Funai

## Abstract

Exercise capacity is a strong predictor of all-cause mortality. Skeletal muscle mitochondrial respiratory capacity, its biggest contributor, adapts robustly to changes in energy demands induced by contractile activity. While transcriptional regulation of mitochondrial enzymes has been extensively studied, there is limited information on how mitochondrial membrane lipids are regulated. Herein, we show that exercise training or muscle disuse alters mitochondrial membrane phospholipids including phosphatidylethanolamine (PE). Addition of PE promoted, whereas removal of PE diminished, mitochondrial respiratory capacity. Surprisingly, skeletal muscle-specific inhibition of mitochondrial-autonomous synthesis of PE caused a respiratory failure due to metabolic insults in the diaphragm muscle. While mitochondrial PE deficiency coincided with increased oxidative stress, neutralization of the latter did not rescue lethality. These findings highlight the previously underappreciated role of mitochondrial membrane phospholipids in dynamically controlling skeletal muscle energetics and function.

## Introduction

Low aerobic capacity is a stronger risk factor for all-cause mortality compared to other common risk factors such as hypertension, type 2 diabetes, and smoking.^1^ Skeletal muscle mitochondrial respiration is the largest contributor for whole-body aerobic capacity,^2^ which in turn is influenced by mitochondrial density and activities of the electron transport system (ETS). Changes in physical activity robustly alters skeletal muscle mitochondrial content and maximal aerobic capacity.^3,4^ Such proliferation or diminishment of mitochondrial biomass must coincide with synthesis or degradation of mitochondrial enzymes and structural lipids. While processes that regulate mitochondrial enzymes are well described,^5,6^ it is unknown how composition of mitochondrial lipids change in response to these adaptations.

Lipids of the inner mitochondrial membrane (IMM) are largely phospholipids with only trace amounts of sphingolipids and cholesterol.^7^ They consist of phosphatidylcholine (PC, 38-45%), phosphatidylethanolamine (PE, 32-39%), cardiolipin (CL, 14-23%), phosphatidylinositol (PI, 2-7%), phosphatidylserine (PS), phosphatidylglycerol (PG), and lyso-phosphatidylcholine (lyso-PC, all less than 3%).^8^ These phospholipids not only give rise to the shape of IMM but are also essential for activities of the enzymes of ETS.^8,9^ In particular, PE and CL are conical-shaped phospholipids that promote the formation of cristae where ETS enzymes reside. These non-bilayer lipids lessen torsional strain of the IMM by localizing into the negatively curved inner leaflet.^10^ They also bind with high affinity to mitochondrial respiratory complexes and regulate their functions.^8,11^ Human mutations that promote loss of mitochondrial PE or CL are detrimental to health.^12-14^

In this study, we set out to describe changes in skeletal muscle mitochondrial phospholipidome that occurs with exercise or disuse. Mitochondrial PE emerged as a key lipid signature that was induced by alterations in physical activity. We then pursued the cellular consequences of changes in muscle mitochondrial PE per se in skeletal muscle-specific tamoxifen-inducible gain- or loss-of-function mouse models. Greater mitochondrial PE, in the absence of changes in the abundance of ETS enzymes, was sufficient to increase the capacity for oxidative phosphorylation. Loss of mitochondrial PE proved fatal due to metabolic and contractile failure in the diaphragm muscle.

## Results

Endurance exercise training induces a robust proliferation of skeletal muscle mitochondria to increase aerobic capacity,^3^ but it is unknown whether training coincides with qualitative changes in mitochondrial phospholipid composition.^8^ C57BL6/J mice were subject to 5-wk of graded treadmill training which promoted skeletal muscle mitochondrial biogenesis (Supplement Figure 1A). Phospholipid analyses of these mitochondria revealed a disproportionately greater increase in PE compared to other phospholipids (Figure 1A&B). High-capacity running (HCR) rats, which had been selectively bred for their intrinsic exercise capacity, demonstrate protection from a wide range of metabolic and cardiovascular diseases compared to low-capacity running (LCR) rats.^15^ Skeletal muscle mitochondria from HCR rats contained more PE than did LCR (Supplement Figure 1B). These observations led us to examine the possibility that an increase in mitochondrial PE contributes to increased aerobic capacity in exercise-trained mice or HCR rats.

**Figure 1.**
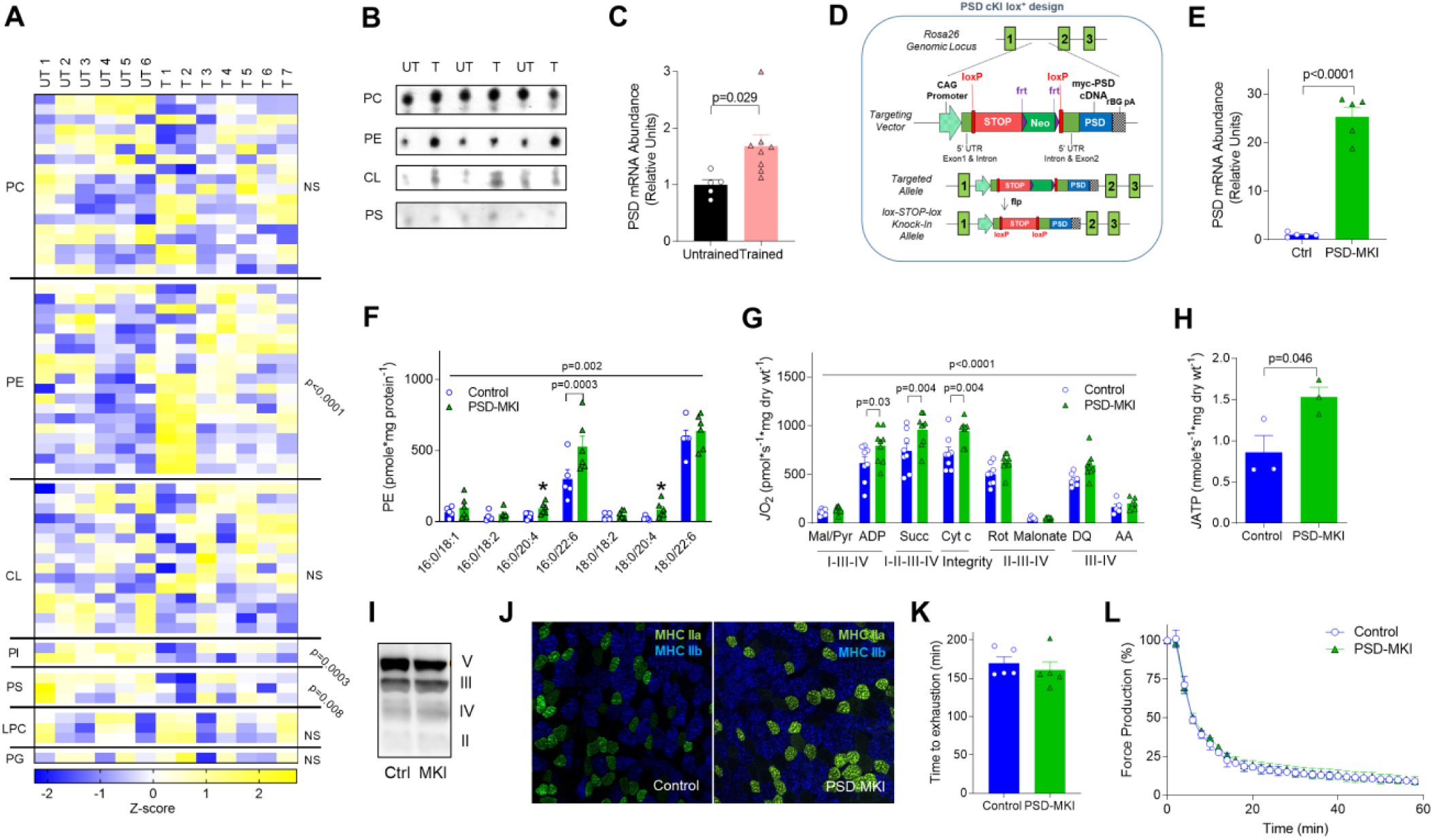
Skeletal muscle mitochondrial PE promotes oxidative capacity. (A-C) Untrained (UT, n=6) or trained (T, n=7 or 8) C57BL6J mice. (A) Skeletal muscle mitochondrial phospholipidome. (B) Mitochondrial phospholipids quantified by TLC. (C) Skeletal muscle PSD mRNA. (D-L) Studies on PSD-MKI mice (n=3-9). (D) Generation of mice with conditional knock-in of PSD. (E) Skeletal muscle PSD mRNA. (F) Muscle mitochondrial PE. (G, H) Rates for oxygen consumption or ATP production in permeabilized muscle fibers with Krebs cycle substrates. (I) Protein abundance of respiratory complex II-V. (J) Myosin-heavy chain fiber-type distribution. (K) Endurance running test. (L) *Ex vivo* twitch endurance test. Mean ±SEM.

Mitochondrial PE is synthesized primarily by the enzyme phosphatidylserine decarboxylase (PSD) that resides in the IMM.^16,17^ Skeletal muscle PSD expression was greater in exercise-trained mice compared to sedentary mice (Figure 1C) and in HCR rats compared to that in LCR rats.^18^ Overexpression of PSD in murine C2C12 myotubes increased the maximal O_2_ consumption rate (Supplement Figure 1C), suggesting that an increased amount of mitochondrial PE enhances respiratory capacity. To study the effects of increased mitochondrial PE *in vivo*, we generated mice with tamoxifen-inducible skeletal-muscle specific overexpression of PSD (PSD-MKI) (Figure 1D). This strategy successfully yielded mice with skeletal muscle-specific PSD overexpression (Figure 1E) and elevated mitochondrial PE (Figure 1F). High-resolution respirometry/fluorometry experiments revealed that PSD overexpression increased the rates of O_2_ consumption and ATP production (Figure 1G&H), effects that were not due to increased mitochondrial mass, abundance of ETS enzymes (Figure 1I, Supplement Figure 1D), or fiber-type (Figure 1J). However, the increase in respiratory capacity did not increase treadmill endurance performance (Figure 1K) or skeletal muscle force generating capacity *ex vivo* (Figure 1L, Supplement Figure 1E&F). PSD-MKI and control mice also did not differ in body weight or composition, food intake, or energy expenditure (Supplement Figure 1G-K). Thus, an increase in muscle mitochondrial PE can increase oxidative capacity but not to an extent that influences endurance. An increase in muscle endurance likely requires concomitant improvements in contractile elements and substrate mobilization.

Skeletal muscle disuse rapidly reduces mitochondrial mass and function. Reduced mitochondrial function precedes disuse-induced muscle atrophy^19^ and might contribute to the mechanism for skeletal muscle loss.^20^ We subjected C57BL6/J mice to a 2-wk hindlimb unloading^21^ which promoted robust muscle loss (Figure 2A). Phospholipid analyses of skeletal muscle mitochondria revealed that disuse promotes an accelerated loss of PE (Figure 2B), concomitant with reduced PSD mRNA (Figure 2C). To model the loss of mitochondrial PE *in vitro*, we performed lentivirus-mediated knockdown of PSD in C2C12 myotubes (Supplement Figure 2A). Reduction in mitochondrial PE (Supplement Figure 2B) robustly reduced mitochondrial respiratory capacity (Supplement Figure 2C&D) in the absence of changes in ETS enzymes (Supplement Figure 2E&F), suggesting that lack of PE suppresses activities of membrane-bound ETS enzymes (Supplement Figure 2G&H). Since global knockout of PSD is embryonically lethal,^22^ we generated mice with tamoxifen-inducible skeletal muscle-specific knockout of PSD (PSD-MKO) (Figure 2D&E). Skeletal muscle mitochondria from PSD-MKO mice were selectively depleted in PE esterified with polyunsaturated fatty acids (Figure 2F&G, Supplement Figure 2I&J). Strikingly, tamoxifen-induced KO promoted a rapid weight loss (Figure 2H), kyphosis (Figure 2I), and ultimately death between 6 and 8-wk after tamoxifen injection (Figure 2J). The lethality of PSD knockout was likely induced by ventilatory failure in respiratory muscles, a common symptom in mitochondrial diseases,^23^ as evidenced by reduced breathing rate and SpO_2_ (Figure 2K&L). Cardiomyopathy, pulmonary edema, low bone density, hypophagia, or hypomobility did not explain the premature death in PSD-MKO mice (Supplement Figure 2K-S). Reduction in body weight was manifested in both lean and fat mass loss (Supplement Figure 2T) as well as in weights of individual muscles including diaphragm (Figure 2M, Supplement Figure 2U). The loss in muscle weights were explained by reduction in cross-sectional area of individual muscle fibers (Figure 2N, Supplement Figure 2V&W). The diaphragm displayed substantial fibrosis (Figure 2O) and loss of force-generating capacity (Figure 2P, Supplement Figure 2X&Y). Thus, acute loss of mitochondrial PE promotes a rapid loss of skeletal muscle mass and function that is reminiscent of atrophy found in disuse in limb muscles as well as that in respiratory muscles during mechanical ventilation.^24^

**Figure 2.**
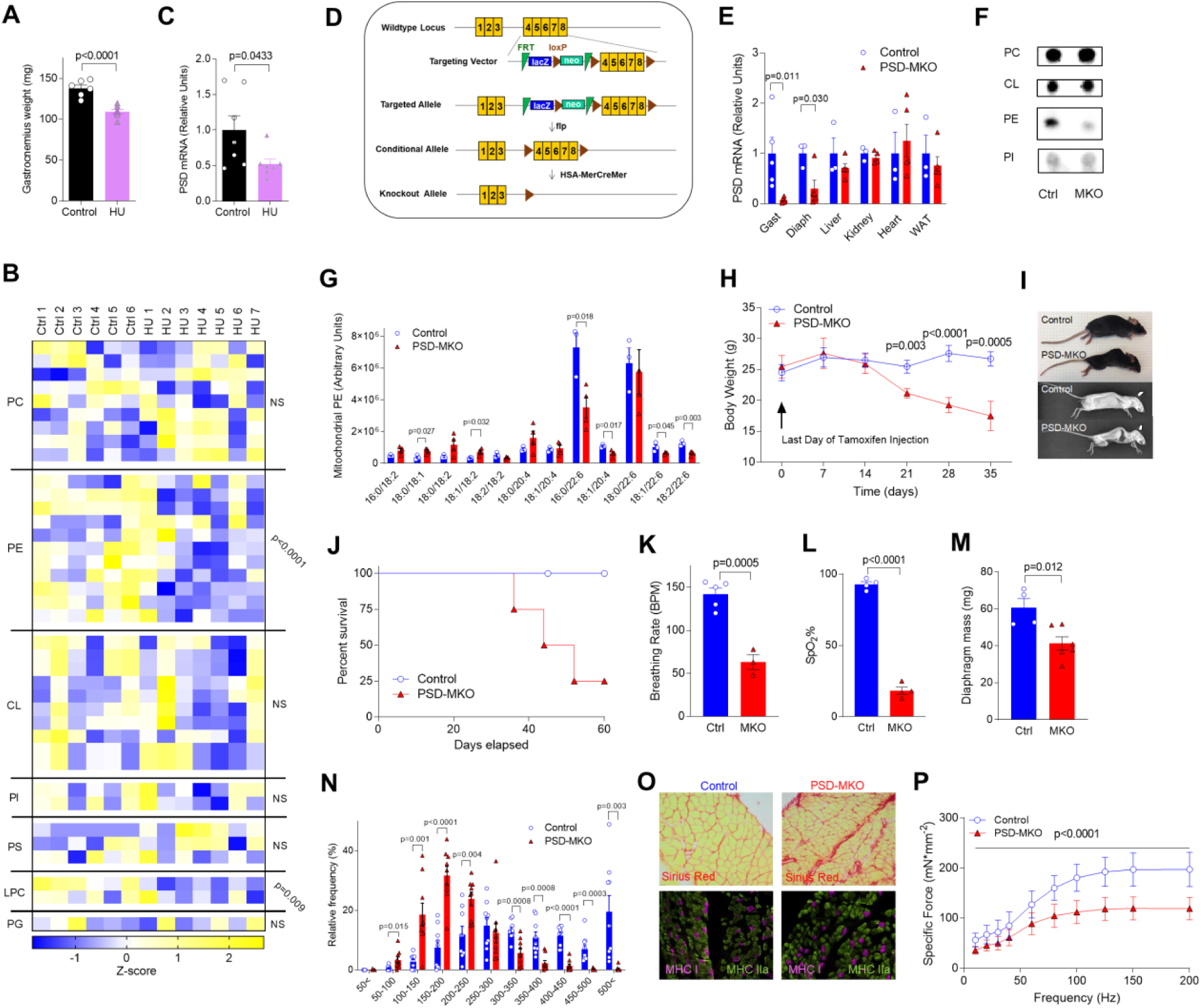
Deficiency of mitochondrial PE promotes atrophy and respiratory failure. (A-C) Control (Ctrl, n=6 or 7) and hindlimb-unloaded (HU, n=7) C57BL6J mice. (A) Gastrocnemius weight. (B) Skeletal muscle mitochondrial phospholipidome. (C) Skeletal muscle PSD mRNA. (D-P) Studies on PSD-MKO mice. (D) Generation of PSD-MKO mice. (E) PSD mRNA levels in multiple tissues (n=5-6). (F) TLC analysis of mitochondrial phospholipids. (G) Muscle mitochondrial PE (n=3-4). (H) Body weights after tamoxifen injection (n=9-22). (I) Kyphosis in PSD-MKO mice. (J) Kaplan-Meier survival curve. (K, L) Breathing rate and peripheral capillary oxygen saturation (SpO_2_) 6-wk post-tamoxifen injection (n=3). (M-P) Diaphragm 4-wk post-tamoxifen injection. (M) Diaphragm weight (n=4-6), (N) distribution of fiber cross-sectional area (n=9), (O) fibrosis and fiber-type, (P) force-frequency curve (n=4-6). Mean ±SEM.

As PSD generates PE for the IMM, the underlying cause of lethal myopathy in PSD-MKO mice is also likely due to changes to mitochondria. PSD deletion deformed mitochondria with less dense cristae (Figure 3A), similar to findings in PSD-depleted CHO cells^25^ and global PSD knockout mice.^22^ These changes occurred in the absence of alteration in abundance of proteins involved in mitochondrial fusion and fission (Supplement Figure 3A). High-resolution respirometry and fluorometry experiments revealed a robust reduction in the rates of O_2_ consumption and ATP production in PSD-MKO muscles (Figure 3B&C, Supplement Figure 3B&C), without changes in abundance of ETS enzymes (Figure 3D). PE molecules are bound to ETS complexes I, II, III, and IV, likely facilitating conformational changes and acting as an allosteric activator.^26-29^ Indeed, enzyme activity assays revealed that activities of ETS complexes I-IV, but not V, were lower in muscles from PSD-MKO mice than in control muscles (Figure 3E). Oxidative phosphorylation is also dependent upon assembly of respiratory supercomplexes,^30,31^ and PE appears to be essential for this process.^25^ Indeed, reduction in mitochondrial PE essentially eliminated the formation of respiratory supercomplexes (Figure 3F). Together, these observations suggest that mitochondrial PE deficiency stagnates efficient electron transfers in the ETS. In turn, inefficiency in electron transfer is predicted to promote electron leakage that causes superoxide production.^32,33^ Indeed, H_2_O_2_ production was markedly higher in PSD-MKO muscles than in control muscles under various substrate conditions (Figure 3G, Supplement Figure 3D-F). Elevated oxidative stress also increased reactive lipid aldehydes such as 4-hydroxynenal and malondeldehyde (Figure 3H&T),^34^ oxidized glutathione (Figure 3J), and counter-oxidative response proteins (Supplement Figure 3G) in PSD-MKO muscles to a greater extent than in control muscles. As oxidative stress has been implicated in skeletal muscle atrophy,^35,36^ we further tested the mechanistic link among mitochondrial PE deficiency, oxidative stress, and respiratory failure in PSD-MKO mice.

**Figure 3.**
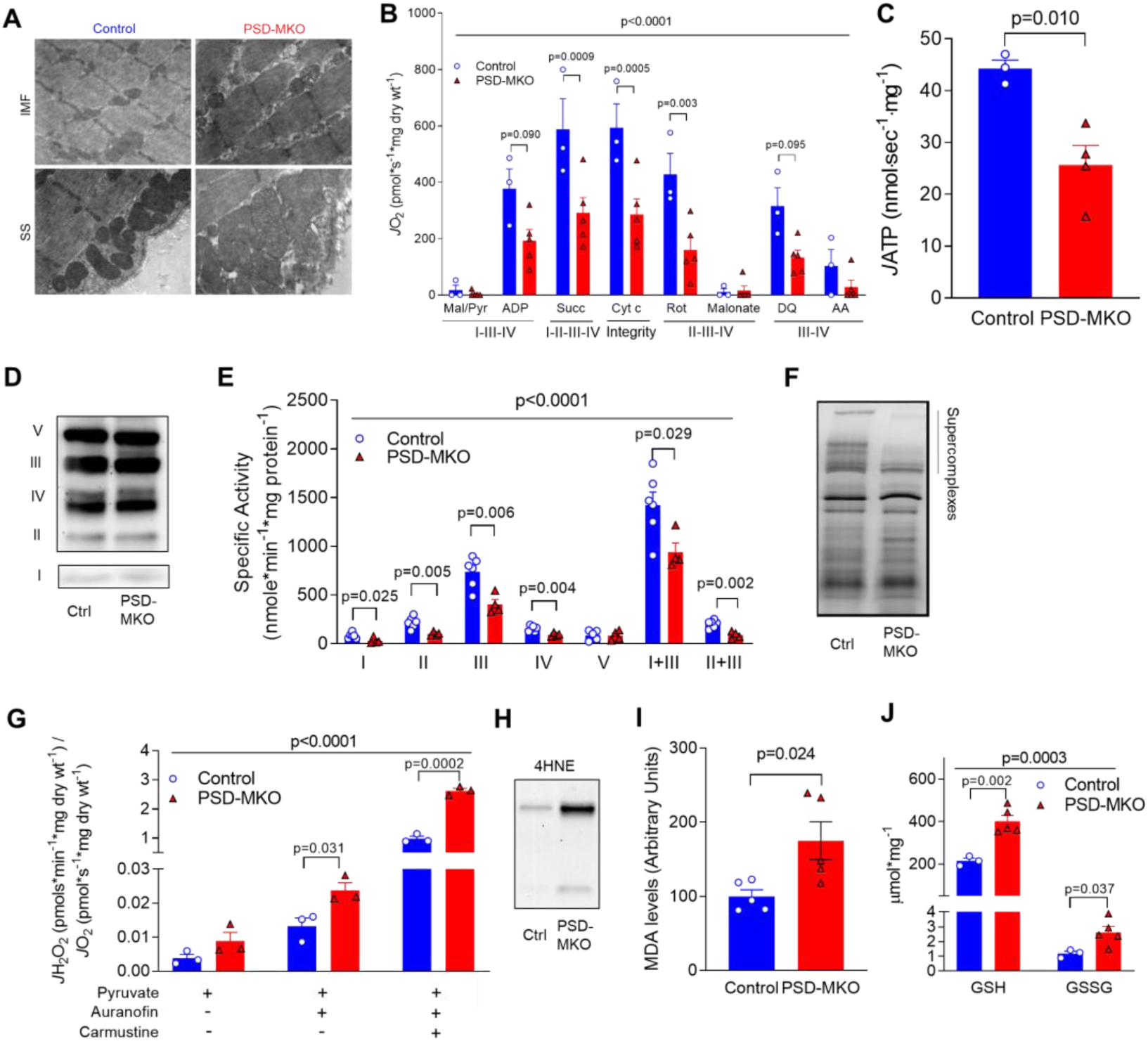
PE deficiency in skeletal muscle mitochondria. (A) Electron micrograph of subsarcolemmal and intermyofibrillar mitochondria. (B, C) Rates of oxygen consumption and ATP production in permeabilized fibers with Krebs cycle substrates (n=3-5). (D) Protein abundance of respiratory complex I-V. (E) Activities of respiratory enzymes (n=4-6). (F) Blue native gel of isolated mitochondria revealing supercomplexes. (G) Mitochondrial H_2_O_2_ production and emission with pyruvate (n=8-9). (H) 4-hydroxy-2-nonenal (4-HNE). (I) malondialdehyde (MDA, n=5). (J) Reduced glutathione (GSH) and oxidized glutathione (GSSG) (n=3-5). Mean ±SEM.

In skeletal muscle, excess superoxide dismutase converts superoxides to H_2_O_2_.^37,38^ To neutralize H_2_O_2_ produced in mitochondrial PE deficient mice, we crossed the PSD-MKO mice with mice that overexpressed mitochondrial-targeted catalase (mCAT) (Figure 4A, Supplement Figure 4A). This strategy yielded mice (mCATxPSD-MKO) with repressed skeletal muscle H_2_O_2_ production (Figure 4B, Supplement Figure 4B). However, mCAT overexpression did not rescue the lethality of PSD-MKO mice (Figure 4C), nor did it ameliorate muscle atrophy (Figure 4D, Supplement Figure 4C), force generating capacity (Figure 4E, Supplement Figure 4D), or oxidative capacity (Figure 4F, Supplement Figure 4E). Thus, while mitochondrial PE deficiency increases oxidative stress, it is not directly responsible for the contractile and metabolic defects in the PSD-MKO mice. This finding was somewhat surprising since oxidative stress is predicted to activate an array of downstream pathways, many of which overlap with defects observed in PSD-MKO mice.^35,36^ To understand the biological processes in PSD-MKO mice that might explain their lethality, we performed deep-sequencing analyses on diaphragms from control, PSD-MKO, and mCATxPSD-MKO mice. Transcripts for 6,026 genes were differentially expressed between control and PSD-MKO diaphragms, and some of which were rescued with mCAT overexpression (Supplement Figure 4F). However, a large majority of differentially-expressed genes were not rescued with mCAT (Supplement Figure 4G), consistent with our observations that deficiency in mitochondrial PE triggers events independent of oxidative stress. Among the 68 pathways that were statistically significantly affected between control and PSD-MKO diaphragms, 45 of them remained altered in mCATxPSD-MKO diaphragms (Figure 4G). Of particular interest, PSD deletion activated pathways for proteasome and ubiquitin-mediated proteolysis, but not lysosome or apoptosis, suggesting that mitochondrial PE deficiency likely promotes muscle atrophy via proteasomal degradation (Figure 4H). Interestingly, PSD deletion also induced activation of transcriptional and translational pathways that were not reversed in mCATxPSD-MKO diaphragms. As expected, mCAT overexpression suppressed activation of antioxidant pathways including glutathione metabolism and peroxisomal genes.

**Figure 4.**
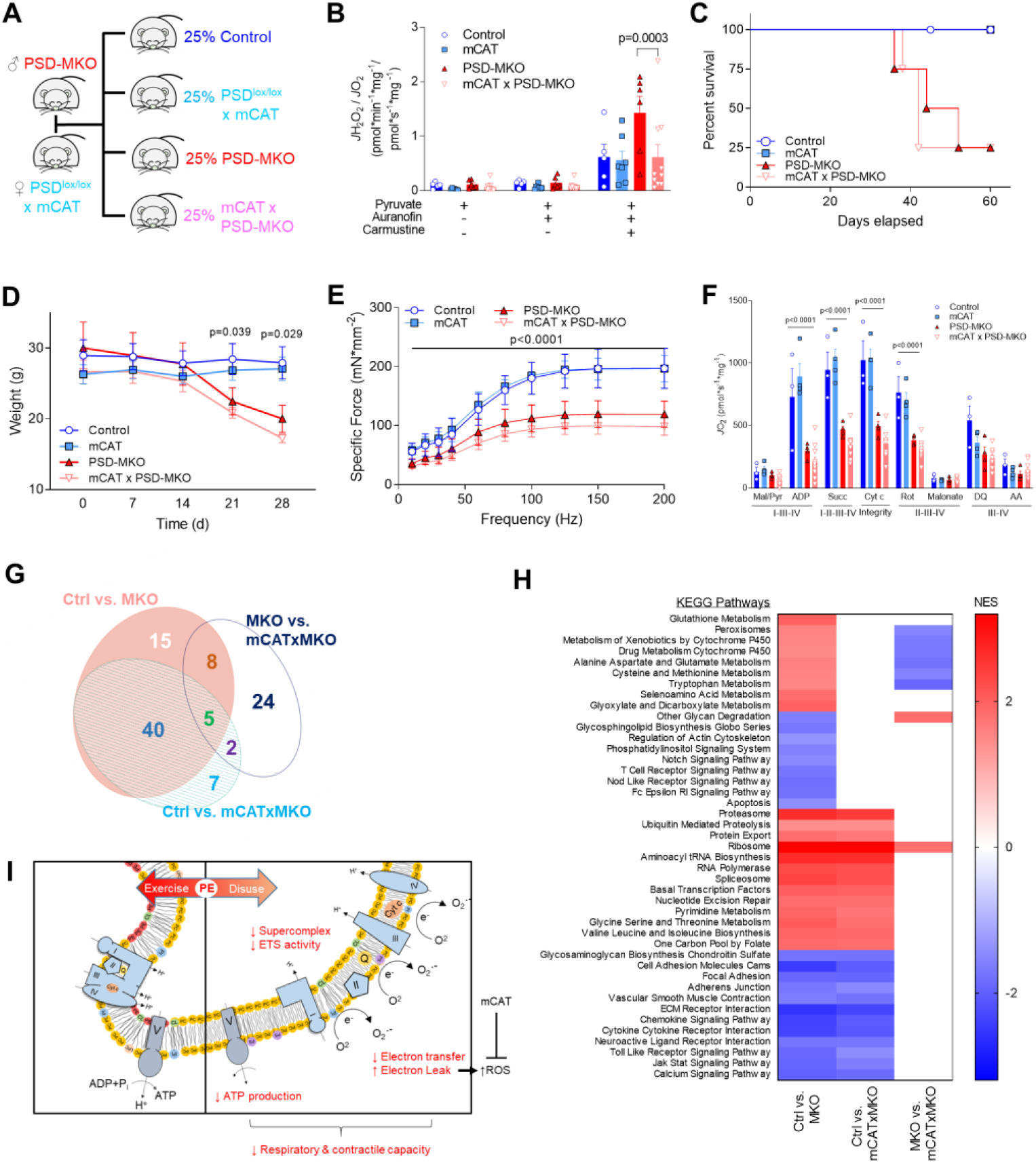
Overexpression of mitochondrial catalase does not rescue PSD deficiency. (A) PSD-MKO mice were crossed with mCAT transgenic mice to generate mCAT x PSD-MKO mice. (B) Mitochondrial H_2_O_2_ production and emission with pyruvate (n=5-7). (C) Kaplan-Meier survival curve. (D) Body weights after tamoxifen injection (n=6-9). (E) Force-frequency curve of diaphragm muscles (n=4-11). (F) Rates of oxygen consumption in permeabilized fibers with Krebs cycle substrates (n=4-10). (G, H) Pathway analyses for differentially-expressed genes between control, PSD-MKO, and mCATxPSD-MKO diaphragms (n=3-4). (G) Area-proportional Venn diagram of differentially-activated pathways. (H) Normalized enrichment scores (NES) of differentially-activated pathways. (I) Schematic illustration of the consequences of mitochondrial PE deficiency. Mean ±SEM.

## Discussion

Phospholipid molecules are largely insoluble in the aqueous cytosol and their intracellular movements are relatively limited. Thus, the membrane phospholipid composition of different organelles is highly distinct, thereby creating a biophysical environment unique to each subcellular location. In mitochondria, the high concentration of PE not only promotes membrane curvature in the cristae, but is also essential for efficient electron transfer and oxidative phosphorylation.^25,39,40^ Recent reports identified loss-of-function mutations in the human *PISD* gene (that encodes PSD enzyme) that promotes severe mitochondrial dysfunction and characterized by congenital cataracts, short stature, facial dysmorphism, platyspondyly, ataxia, and/or intellectual disability.^12,14^ Combined with data presented in the current manuscript, these findings indicate a critical role that mitochondrial PE plays in health and disease.

In skeletal muscle, inhibition of mitochondria-autonomous synthesis of PE via PSD causes robust skeletal muscle atrophy and ventilatory failure due to fibrosis and loss of contractility in the diaphragm muscle. The underlying cause appears to be due to a system failure of mitochondrial circuitry that is evident in reduced activities of complex I-IV, low supercomplex formation, and elevated electron leak (Figure 4I). We predict that these defects trigger proteosomal degradation pathways that promote muscle atrophy and weakness. In contrast, such a phenotype does not occur when PE synthesis via the CDP-ethanolamine pathway on the ER is inhibited in skeletal muscle,^41,42^ suggesting that pools of PE made by PSD and CDP-ethanolamine pathway are functionally distinct. How select phospholipids (such as PC, PI and PA) are transported from ER to mitochondria, but PE is unable to do so is not understood.^17,40^ Furthermore, it is known that PE generated by PSD is readily transported from mitochondria to ER.^43^ Thus it remains possible that some of the defects in the PSD-MKO mice are due to a lack of PE exported from mitochondria.

In conclusion, our findings reveal that alteration in PE composition represents a key adaptive response to exercise or disuse in skeletal muscle mitochondria. Gain- or loss-of-function studies show that changes in mitochondrial PE modulates oxidative capacity independent of changes in abundance of ETS proteins. A deficiency of mitochondrial PE in the diaphragm muscle is detrimental to its metabolic and contractile function, leading to ventilatory failure and lethality. While these defects were associated with increased oxidative stress, its neutralization with mitochondrial-targeted catalase did not prevent any of the dysfunction induced by PSD deficiency. These findings also raise a possibility that reduced mitochondrial PE represents a mechanism by which disuse promotes the loss of muscle mass and function associated with muscle atrophy. Changes in mitochondrial phospholipid composition appears to be an important regulatory mechanism by which physical activity modulates mitochondrial energetics in skeletal muscle.

## Materials and Methods

### Rodent models

PSD conditional knock-in (PSDcKI^+/+^) mice were generated by inserting myc-tagged mouse *Pisd* cDNA into a Rosa26 locus. *Pisd* cDNA was preceded by a CAG promoter and loxP-flanked stop codon for a tissue-specific ectopic expression of PSD. These mice were then crossed with HSA-MerCreMer mice (tamoxifen-inducible α-human skeletal actin Cre, courtesy of Dr. Karyn Esser, University of Florida) to generate PSD-MKI (PSDcKI^+/-^, HSA-MerCreMer^+/-^) and control (PSDcKI^+/-^, No Cre) mice. Mouse embryonic stem (ES) cells that carried loxP sites flanking exons 4-8 of the mouse PSD gene were purchased from the European Conditional Mouse Mutagenesis Program (EUCOMM). The cells were microinjected into C57BL/6J blastocysts and transplanted in pseudopregnant females to produce PSD conditional knockout (PSDcKO^+/+^) mice. These mice were crossed with HSA-MerCreMer mice to generate PSD-MKO (PSDcKO^+/+^, HSA-MerCreMer+/-) and control (PSDcKO^+/+^, No Cre) mice. Mitochondrial catalase (mCAT) transgenic mice were purchased from the Jackson Laboratory (Stock No: 0161971) and crossed with PSD-MKO mice to generate the mCATxPSD-MKO (mCAT^+/-^, PSDcKO^+/+^, HSA-MerCreMer^+/-^), PSD-MKO, mCAT (mCAT^+/-^, PSDcKO^+/+^, No Cre), and control (PSDcKO^+/+^, no Cre) mice. Cre control mice (PSDcKO^-/-^ or PSDcKI^-/-^, HSA-MerCreMer^+/-^) and tamoxifen-untreated control mice displayed no difference in phenotype to loxP control mice. All mice were bred onto C57BL/6J background and were born at normal Mendelian ratios. Both male and female mice were studied with no difference in phenotypes. HCR and LCR rats were maintained and studied at the University of Michigan. All animals were fasted 4 h prior to tissue collection. All protocols were approved by Institutional Animal Care and Use Committees at University of Utah, East Carolina University, and University of Michigan.

### Exercise training

Male C57BL/6J mice were kept untrained (n = 5) or underwent treadmill training (5 d/wk, 12 m/min, 2-6% incline, n = 8) for 5 weeks. The mice were then sacrificed and tissues were dissected ∼40 h after the last exercise session.

### Hindlimb unloading

Male C57BL/6J mice underwent 2 weeks of hindlimb unloading (HU) or were ambulatory controls. The HU (2 mice/cage) were subjected to a modified unloading method based on the traditional Morey-Holton design for studying disuse atrophy in rodents.^21^ Body weight and food intake were monitored every other day to ensure that mice did not experience excessive weight loss due to malnutrition or dehydration. At the end of day 14 of HU, mice were fasted for 4 h and anesthetized for tissue collection.

### RNA quantification

For qPCR experiments, mouse tissues or cells were lysed in 1 ml of Trizol (ThermoFisher) and RNA was isolated using standard techniques. The iScript™ cDNA synthesis kit was used to reverse transcribe total RNA and quantitative PCR was performed with SYBR Green® reagents (ThermoFisher). Pre-validated primer sequences were obtained from mouse primer depot (https://mouseprimerdepot.nci.nih.gov/). All mRNA levels were normalized to RPL32. For RNA sequencing, diaphragm RNA was isolated with RNeasy Kit (Qiagen, #74104). RNA library construction and sequencing were performed by the High-Throughput Genomics Core at the Huntsman Cancer Institute/University of Utah. RNA libraries were constructed using the Illumina TruSeq Stranded Total RNA Sample Prep Kit and contaminating rRNAs were removed using RiboZero Gold. Sequencing was performed using a NovSeq2 with 25 million reads per sample. Pathway analyses were performed by the Bioinformatics Core at the Huntsman Cancer Institute/University of Utah using the KEGG Pathway database. For differentially-expressed genes, only transcripts with P_adj_ < 0.05 and BaseMean > 100 are included. For pathway analyses, the area-proportional Venn diagram was drawn with eulerAPE.^44^ KEGG Pathways that were differentially expressed between control and PSD-MKO diaphragms were stratified to those that were or were not rescued by mCAT overexpression.

### Mitochondrial isolation

Tissues were minced in ice cold MIM buffer (300 mM Sucrose, 10 mM HEPES, 1 mM EGTA, 1 mg/ml BSA, pH 7.4) and gently homogenized with a Teflon pestle. The homogenate was centrifuged at 800 × g for 10 min at 4°C. The supernatant was transferred to another tube and centrifuged again at 12,000 × g for 10 min at 4°C. The crude mitochondrial pellet was suspended in 15% Percoll (diluted with MIM buffer) and a discontinuous Percoll gradient was prepared consisting of 50%, 22%, and 15% Percoll layers. Mitochondria were carefully layered on top of the gradient and spun at 22,700 RPM for 10 min at 4°C in an ultracentrifuge (Thermo Scientific SureSpin 630 rotor). The purified mitochondrial fraction was collected at the 50% - 22% Percoll interface. To remove excess Percoll, the collected mitochondrial fraction was diluted with MIM and spun for 3 min at 10,000 ×g. This step was repeated twice and the final mitochondrial pellet was suspended in MIM buffer for experiments.

### Lipid extraction, thin layer chromatography, and lipid mass spectrometry

Mitochondrial lipids were extracted using a modified Bligh-Dyer extraction. Resuspended lipids were then used for phospholipid quantification by thin layer chromatography or mass spectrometry. The TLC plates were developed using chloroform:glacial acetic acid:methanol:water (65:35:5:2) as mobile phase for PC and PS, and chloroform:glacial acetic acid:methanol:water (85:25:5:2) as mobile phase for PE, CL, and PI. The plates were dried, sprayed with charring solution, and heated at 190 °C for ∼15 min. Intensity of the charred lipid spots was measured using an Odyssey Infrarred Imager. Mass spectrometry analyses of phospholipids for exercise training, disuse, and PSD-MKI samples were performed at the University of Utah Metabolomics Core, with untargeted (Agilent 6530 UPLC-QToF mass spectrometer) and targeted (UPLC-QQQ mass spectrometer) platforms. Mass spectrometric analyses of HCR/LCR and PSD-MKO samples were performed at the University of Michigan Nutrition and Obesity Research Center Metabolomics Core using a ABSCIEX 5600 TripleTOF mass spectrometer. For untargeted comprehensive lipidomics (exercise training, HCR/LCR, disuse), quantities are expressed as z-scores.

### Cell culture

C2C12 myoblasts were grown and maintained in high glucose DMEM + 10% fetal bovine serum (FBS) + 100 µg/ml of penicillin/streptomycin. Once 90-100% confluent, C2C12 myoblasts were differentiated into myotubes using low glucose DMEM (1 g/L glucose, L-glutamine, 110 mg/L sodium pyruvate) + 2% horse serum + 100 µg/ml of penicillin/streptomycin. HEK 293T cells were maintained in high glucose DMEM + 10% FBS + 100 µg/ml of penicillin/streptomycin. The overexpression or lentivirus-mediated knockdown of PSD was performed as previously described ^45^. Vectors were sourced from OriGene (Rockville, MD) for PISD-expressing plasmid (MR206380), Sigma (St. Louis, MO) for shRNA for mouse PISD (shPSD: TRCN0000115415, and Addgene (Cambridge, MA) for psPAX2 (ID #12260), pMD2.G (ID #12259), and scrambled shRNA plasmid (SC: ID #1864).

### Mitochondrial respiration measurements

Respiration in permeabilized muscle fiber bundles and isolated mitochondria was performed as previously described.^46,47^ Briefly, a small portion of freshly dissected red gastrocnemius muscle tissue was placed in Buffer X (7.23 mM K_2_EGTA, 2.77 mM Ca K_2_EGTA, 20 mM imidazole, 20 mM taurine, 5.7 mM ATP, 14.3 mM phosphocreatine, 6.56 mM MgCl_2_.6H_2_O, and 50 mM K-MES, pH = 7.1), Fiber bundles were separated and permeabilized for 30 min at 4°C with saponin (30 µg/ml) and immediately washed in Buffer Z (105 mM K-MES, 30 mM KCl, 10 mM K_2_HPO_4_, 5 mM MgCl_2_.6H_2_O, 0.5 mg/ml BSA, and 1 mM EGTA, pH = 7.4) for 15 min. After washing, high-resolution respiration rates were measured using an OROBOROS Oxygraph-2k. The muscle fibers were suspended in Buffer Z with 20 mM creatine and 10 µM blebbistatin to inhibit myosin ATPases during respiration measurements. A variety of respiration protocols were utilized in permeabilized fibers and isolated mitochondria. For the mixed substrate protocol, the chamber was hyperoxygenated to ∼300 pmol and started with the addition of malate (0.5 mM) followed by sequential additions of pyruvate (5 mM), ADP (2 mM), succinate (10 mM), cytochrome c (10 µM), rotenone (5 µM), malonate (5mM), duroquinol (0.25 mM), and antimycin A (2 µM). For the fatty acid oxidation protocol, the chamber was not hyperoxygenated and the protocol started with the addition of malate (0.5 mM) followed by sequential additions of palmitoyl-l-carnitine (50 µM) and ADP (2 mM). For Complex IV mediated respiration, rotenone (5 µM), malonate (5mM), and antimycin A (2 µM) were added to inhibit Complexes I-III, after which ascorbate (2mM, prevents autooxidation of TMPD), TMPD (0.5 mM), and KCN (20 mM) were sequentially added. When respiration experiments were complete, fiber bundles were washed in distilled H_2_O to remove salts, then freeze dried in a lyophilizer (Lab-Conco). Dry weight was measured and respiration rate was expressed relative to fiber weight. When isolated mitochondria respiration rates were measured, the respiration rate was normalized to the total protein content in the chamber.

Oxygen consumption rates in C2C12 cells were measured with a Seahorse Flux Analyzer XF24 or XFe96 (Seahorse Bioscience, Billerica, MA). The cells were plated at ∼20,000-40,000 cells/well and then differentiated into myotubes. On the day of the experiment, medium was switched to XF Assay Medium Modified DMEM (pH = 7.4) containing added glucose (10 mM), pyruvate (200 mM), and glutamine (200 mM) for 1 h. Subsequently, basal and maximal respiration rates were measured as previously described.^46^

### H_2_O_2_ emission and production

The Amplex Ultra Red (10 µM) / horseradish peroxidase (3 U/ml) detection system was used to measure mitochondrial H_2_O_2_ emission and production fluorometrically (Ex:Em 565:600, HORIBA Jobin Yvon Fluorolog) at 37°C.^47^ Permeabilized muscle fibers were placed into a glass cuvette with Amplex Ultra Red reagents and buffer Z (with 1 mM EGTA and 23 U superoxide dismutase). Initially, an 8-min background rate was obtained, followed by addition of palmitoyl-L-carnitine / malate (50 µM / 1 mM) into the cuvette for measurement of H_2_O_2_ emission rate. For maximal H_2_O_2_ production rate, auranofin (1 µM) and carmustine (BCNU, 100 µM) were titrated into the cuvette to inhibit thioredoxin reductase and glutathione reductase, respectively. The fiber bundles were washed in distilled H_2_O, freeze dried, and dry weight was measured. H_2_O_2_ rates were expressed relative to fiber weight. Rates were then corrected for O_2_ consumption, which was measured with an Oxygraph-2k machine in the presence of identical substrates.

### Western blotting

Tissues or cells were homogenized in lysis buffer, nutated at 4°C for 1 h, centrifuged at 4°C for 15 min at 12,000 × g, and the supernatant was transferred to a new tube. Western blotting was performed as previously described (10) and samples were analyzed for protein abundance of FoxO1 (Cell Signaling, #2880), 4-HNE (Abcam, ab48506), DRP-1 (Abcam, ab56788), pDRP-1 (Cell Signaling, #3455), Mfn-2 (Abcam, ab56889), Nrf2 (DSHB), FoxO3 (DSHB), PRDX4 (DSHB), Citrate Synthase (Abcam, ab96600), and catalase (Abcam, ab1877).

### Specific activity of mitochondrial enzymes

The specific activities of electron transport chain enzymes were determined using spectrophotometric methods. Briefly, myoblasts or isolated mitochondria were freeze thawed 2-3 times to disrupt the outer mitochondrial membrane and allow access of substrates to enzymes. The specific activity of each complex was measured at 37°C using reagents and substrates specified previously.^48^ For experiments involving C2C12 cells, Complexes I and V activities were measured in isolated mitochondria, whereas activities of other Complexes were measured in whole cells. Citrate synthase activity was measured on a 96-well plate using a commercially available kit (Sigma Aldrich, CS0720).

### Blue native PAGE

Isolated mitochondria suspended in MIM were solubilized (0.5-2 mg) in 2% digitonin for 15 min on ice and then centrifuged at 20,000 × g for 30 min at 4°C. The supernatant was collected and placed into a new tube, and protein content was measured. Approximately 25-35 µg of mitochondrial protein was suspended in a mix of native PAGE 5% G-250 sample buffer and 1X native-PAGE sample buffer (total volume of 10-20 µl). The samples and standards were then loaded onto a native PAGE 3-12% Bis-Tris Gel (ThermoFisher, BN1001BOX) and electrophoresis performed at 150 V for 3 h on ice. The gel was then placed in fixative solution (40% methanol, 10% acetic acid), microwaved on high for 45 s, and then shaken on an orbital shaker for 15 min at room temperature. After incubation, the gel was placed in destaining solution (8% acetic acid), microwaved on high for 45 s, then incubated overnight at 4°C on an orbital shaker. The gel was scanned for densitometry.

### Glutathione protein abundance

Protein levels of reduced glutathione (GSH) and oxidized glutathione disulfide (GSSG) were measured using high-performance liquid chromatography (Shimadzu Prominence HPLC system). Freshly dissected gastrocnemius muscle was hand homogenized with a glass pestle in buffer containing 50 mmol/L Trizma base, 20 mmol/L boric acid, 20 mmol/L L-serine, and 10 mmol/L N-ethylmaleimide. The homogenate was then frozen until processing, in which the sample was split into two parts for detection of GSH and GSSG. For GSH measurement the homogenate was deproteinated with 1:10 (v/v) 15% trichloroacetic acid, centrifuged for 5 min at 20,000 × *g*, and the supernatant was transferred to an autosampler vial for measurement of GSH by high-performance liquid chromatography using 92.5% of a 0.25% (v/v) glacial acetic acid mixed with 7.5% pure HPLC-grade acetonitrile. Ultraviolet chromatography was used to measure the GSH–NEM conjugate at a wavelength of 265 nmol/L (SPD-20A; Shimadzu).

### Malondialdehyde (MDA) quantification

MDA content was quantified in fresh gastrocnemius muscles using a lipid peroxidation assay kit (Abcam, ab118970) according to manufacturer’s instruction. Rates of appearance of MDA-thiobarbituric acid (TBA) adduct were quantified colorimetrically at 532 nm using a spectrophotometer.

### Bone density, body composition, and indirect calorimetry

Bone density of the femur was determined using microcomputed tomography (µCT, Scanco Medical AG). Fat and lean body mass were measured using the EchoMRI-500 (EchoMRI, Houston, TX). Whole body oxygen consumption, respiratory exchange ratio, and physical activity levels were determined using a TSE LabMaster System (TSE Systems, Chesterfield, MO). Measurements were taken over a 4-5 d period, with the first 1-2 d of data excluded for acclimitization to the new environment. Data represent averages of two or three 12-h light or dark cycles.

### Echocardiography

Echocardiographic measurements were made as previously described.^47^ Briefly, mice were anesthetized with a 0.5-2% isoflurane in an oxygen mixture and kept on a heated monitoring plate to maintain body temperature. Heart rate was kept between 400-500 bpm for all measurements to ensure physiological relevance. The Vevo 2100™ High-Resolution In Vivo Imaging System (VisualSonics) was used with a 30-MHz transducer for echocardiographic recordings. B-mode recordings from transthoracic long-axis view were used to measure left ventricular volume during diastole and systole. These measurements were used to calculate ejection fraction, stroke volume, and cardiac output.

### Electron microscopy

Freshly dissected skeletal muscle was immediately placed in a 2% glutaraldehyde and 0.1 M cacodylate fixation mix and cut into longitudinal sections ∼2 mm in diameter and ∼3-4 mm in length. Tissues were stored in fixation mixture at 4°C until all tissues were collected. The tissues were washed three times for 10 min in 0.1 M phosphate buffer, fixed in 1% oxmium tetroxide for 1 h, then washed three times for 10 min in 0.1 M phosphate buffer. The fixed tissues were dehydrated in sequential steps of 25%, 50%, 75%, and 100% alcohol (twice each step) for 15 min each. The tissue was then embedded in increasing concentrations of Spurrs media including 30% (for 30 min), 70% (overnight), 100% (2 h), and 100% (30 min). Each tissue was placed in a flat-bottom embedding mold filled 50% with Spurrs medium, then polymerized at ∼60°C overnight. The tissues were then cut through the transverse plane and imaged with a JEOL 1200EX transmission electron microscope equipped with a Soft Imaging Systems MegaView III CCD camera.

### Muscle strength experiments

The extensor digitorum longus (EDL) and diaphragm muscles were dissected as previously described^49^ and tied into the Horizontal Tissue Bath System from Aurora Scientific, Inc (Model: 801C). Muscles were stimulated with a 20V twitch train and stretched until optimal length for force production was reached. After a 5 min equilibration period, muscles were stimulated with frequencies ranging from 10-200 Hz (0.1 ms pulse, 330 ms train, 2 min between trains). Forces produced by electrically-stimulated muscle contractions were recorded in real time via a force transducer (Aurora Scientific Inc., Model: 400A). Specific force was calculated using cross sectional area of the muscle tissue (mN/mm^2^), as estimated from the weight and length of the muscle.

### Histology

Frozen muscle or lung tissues were embedded in optimal cutting temperature (OCT) compound and were sectioned (10μm) with a cryostat (Microtome Plus). Muscle sections were used for myosin heavy chain (MHC) isoform immunofluorescence (IF) and Sirius Red staining to examine fibrosis. For MHC IF, sections were incubated with MHCI (BA.D5), MHCIIa (SC.71), MHCIIb (BF.F3, all three from Developmental Studies Hybridoma Bank, University of Iowa), or laminin (Sigma, #L9393) and imaged at the University of Utah Cell Imaging Core. Negative-stained fibers were considered to be MHCIIx. Master Tech Picro Sirius Red was used for Sirius Red staining. Myofiber cross-sectional area was quantified using semi-automatic muscle analysis using segmentation of histology: a MATLAB application (SMASH) alongside ImageJ software. Lung tissues were used for H&E staining.

### Statistical Analyses

Data are presented as means ±SEM. Statistical analyses were performed using GraphPad Prism 7.03 software. Independent samples t-tests were used to compare two groups. For two-by-two comparisons, two-way ANOVA analyses were performed (main effect of genotype shown over a horizontal line) followed by appropriate post-hoc tests corrected for multiple comparisons. All tests were two-sided and p<0.05 was considered statistically significant.

## Acknowledgements

This work was supported by NIH grants DK107397, DK109888, DK095774 (to K.F.), DK110656 (to P.D.N.), AG050781 (to M.J.D.), HL123647, AT008375 (to S.R.S.), HL129632 (T.E.R.), and DK109556 (to T.D.H.), Larry H. & Gail Miller Foundation grant (to P.J.F.), and American Heart Association grants 19PRE34380991 (to J.M.J), 18PRE33960491 (to A.R.P.V.), and 16POST30980047 (to T.D.H.). University of Utah Metabolomics Core Facility is supported by S10 OD016232, S10 OD021505, and U54 DK110858. Mouse lines are available from K.F. upon request.

---

**Supplement Figure 1.**
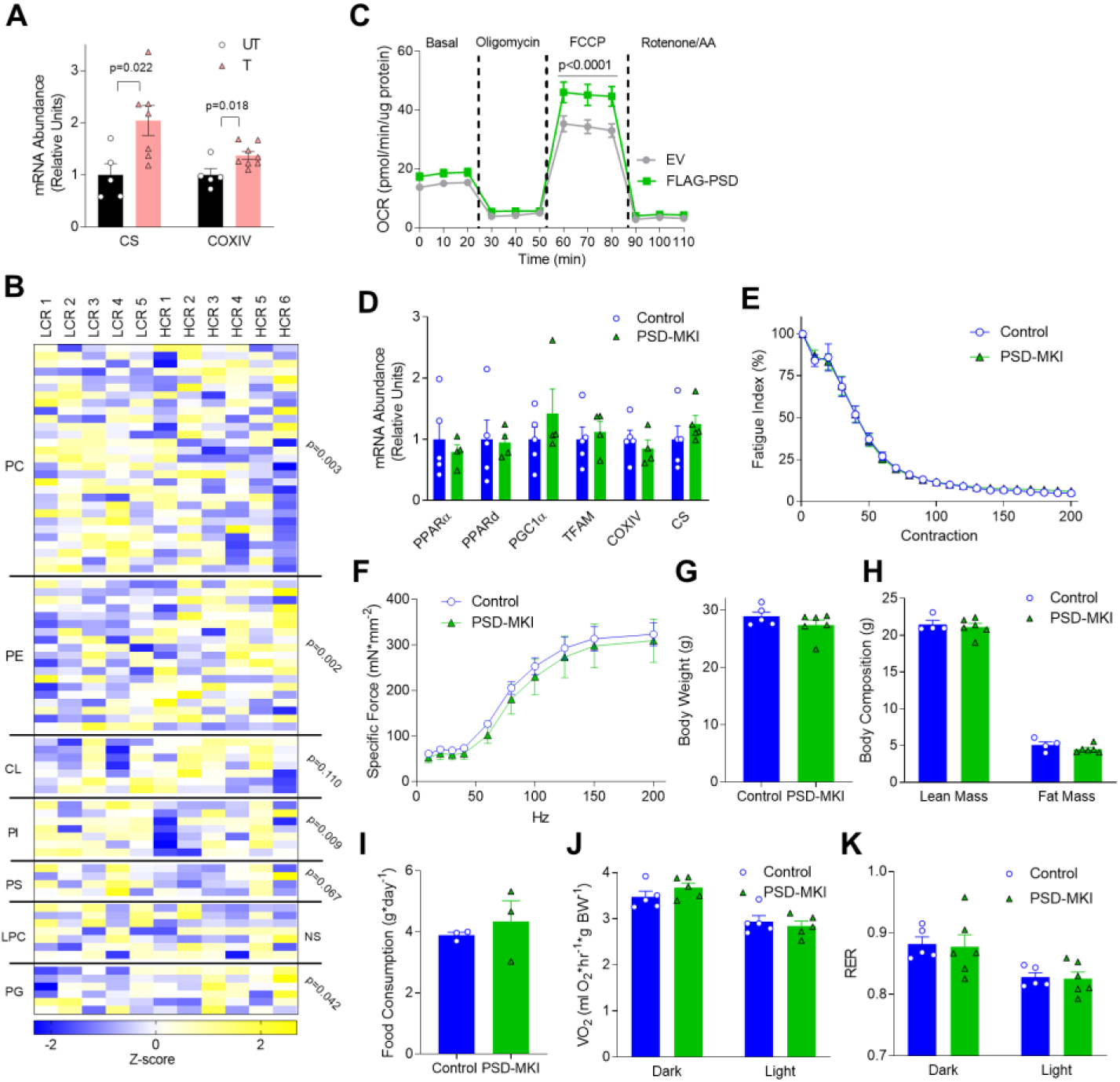
Skeletal muscle mitochondrial PE and oxidative capacity. (A) Skeletal muscle citrate synthase (CS) and cytochrome oxidase IV (COXIV) mRNA in UT or T mice (n=5-8). (B) Skeletal muscle mitochondrial phospholipidome in low-capacity running (LCR) or high-capacity running (HCR) rats (n=5-6). (C) Oxygen consumption rates (OCR) from empty vector (EV) or PSD-expression vector (FLAG-PSD) treated C2C12 myotubes (n=6). (D-K) Ctrl or PSD-MKI mice. (D) Muscle mRNA encoding mitochondrial enzymes and transcription factors (n=5). (E) Fatigue curve with tetanic contractions (n=5-6). (F) Force-frequency curve (n=3). (G) Body weights (n=5-6). (H) Body composition (n=4-6). (I) Food consumption (n=3). (J) Whole-body VO_2_ (n=5). (K) Respiratory exchange ratio (RER, n=5). Mean ±SEM.

**Supplement Figure 2.**
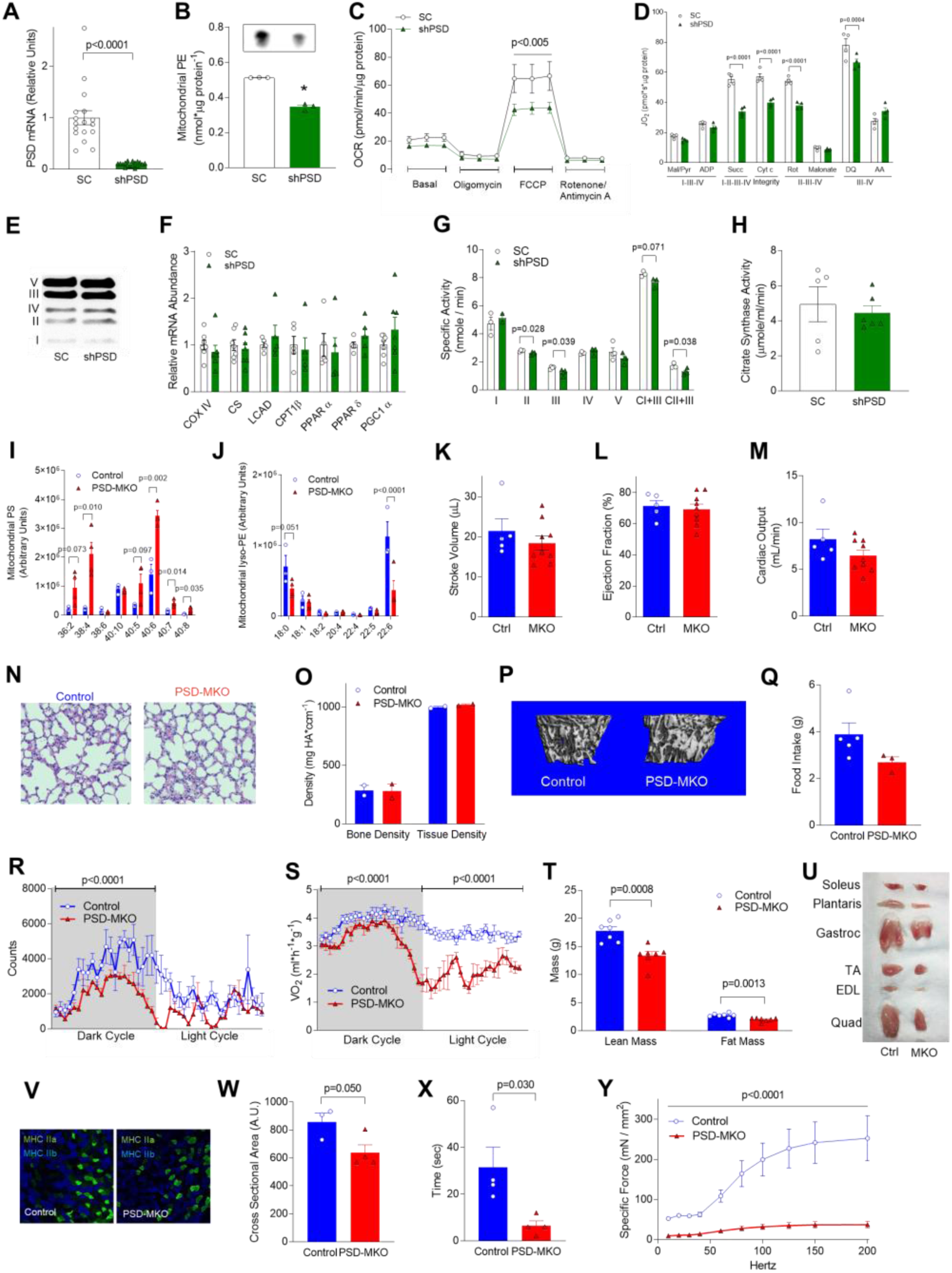
Deficiency of muscle mitochondrial PE *in vitro* and *in vivo*. (A-H) C2C12 myotubes incubated with lentivirus expressing scrambled (SC) or shRNA targeted to PSD (shPSD). (A) PSD mRNA (n=17). (B) Mitochondrial PE (n=3). (C) OCR in intact myotubes (n=6). (D) Rate of oxygen consumption using Krebs cycle substrates (n=4). (E) Protein abundance of respiratory complexes I-V. (F) mRNA encoding mitochondrial enzymes and transcription factors (n=4). (G) Activities of respiratory enzymes (n=3-4). (H) Citrate synthase activity (n=5-6). (I-Y). Studies on PSD-MKO mice. (I) Muscle mitochondrial lyso-PE (n=3-4). (J) Muscle mitochondrial PS (n=3-4). (K-M) Stroke volume, ejection fraction, or cardiac output measured with echocardiography (n=5-8). (N) H&E staining of lung section. (O, P) Bone density by μCT scan (n=2). (Q) Food intake (n=3-5). (R, S) Activity and VO_2_ measured by indirect calorimetry (n=6). (T) Body composition (n=7). (U) Muscle sizes. (V) Fiber-type composition. (W) Fiber cross-sectional area (n=3-4). (X) Kondziela’s inverted screen test (n=4). (Y) Force-frequency curve of extensor digitorum longus muscles (n=2-4). Mean ±SEM.

**Supplement Figure 3.**
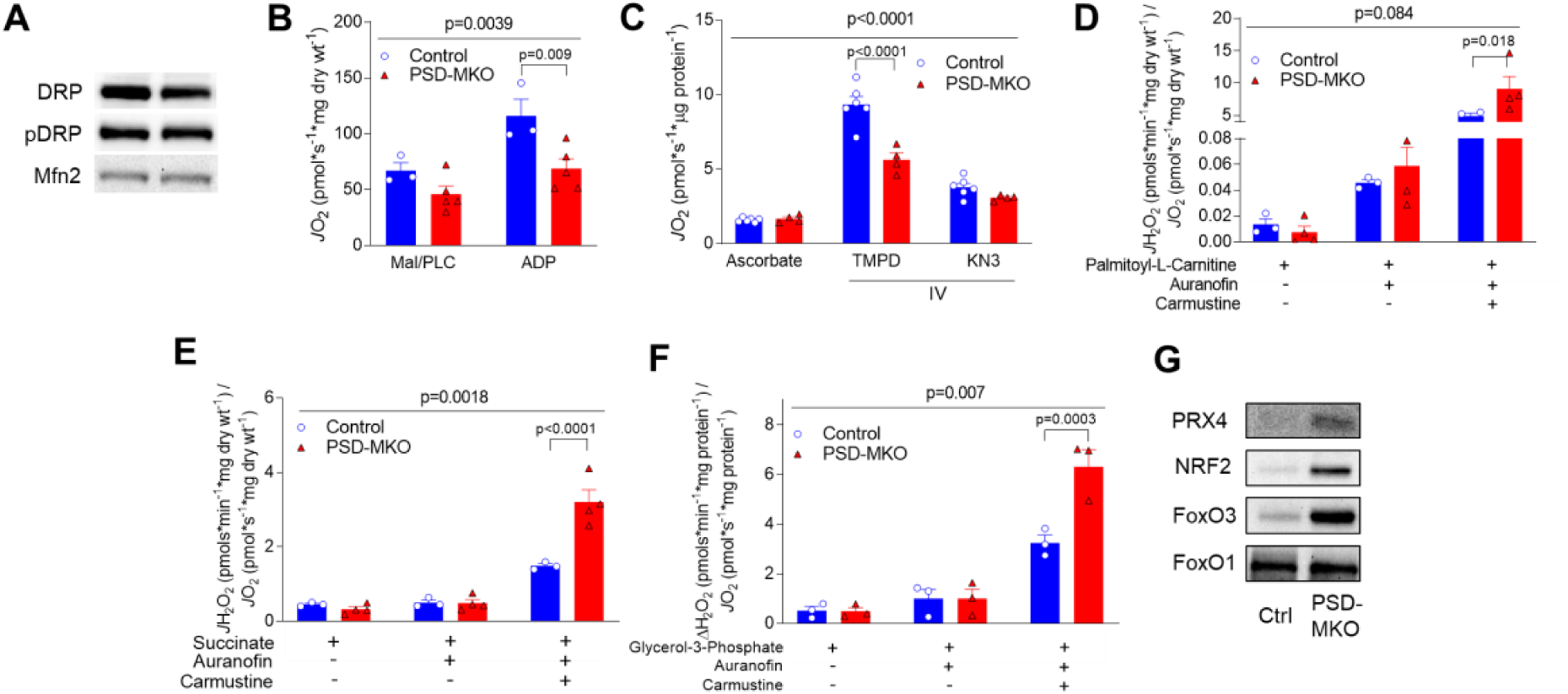
PE deficiency in skeletal muscle mitochondria. (A) Total and phosphorylated (Ser 616) dynamin-related protein (DRP) and mitofusion 2. (B) Palmitoyl-L-carnitine (PLC)-induced oxygen consumption in permeabilized fibers (n=3-5). (C) Complex IV-mediated respiration rates in isolated mitochondria (n=4-6). (D-F) Mitochondrial H_2_O_2_ production and emission with palmitoyl-L-carnitine, succinate, or glycerol-3-phosphate (n=3-4). (G) Protein abundance of the antioxidant enzyme peroxiredoxin 4 (PRX4) and regulators of antioxidant defense including nuclear factor erythroid 2-related factor 2 (NRF2), Forkhead box protein O1 (FoxO1), and FoxO3. Mean ±SEM.

**Supplement Figure 4.**
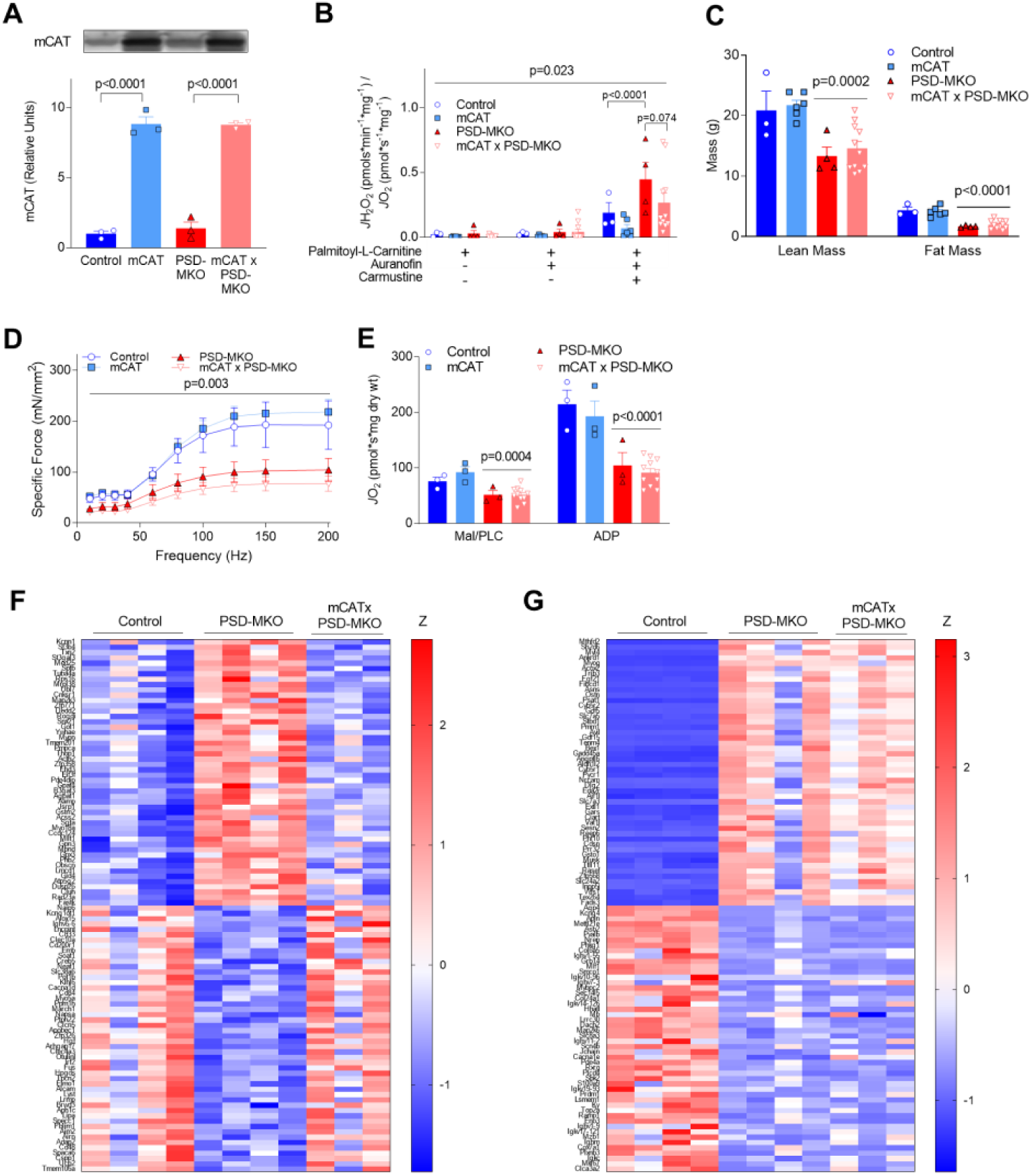
Overexpression of mitochondrial catalase does not rescue muscle-specific PSD deficiency. (A) Protein abundance of mCAT (n=3). (B) Mitochondrial H_2_O_2_ production and emission with palmitoyl-L-carnitine (n=3-11). (C) Body composition of mice 4-wk post-tamoxifen injection (n=3-11). (D) Force frequency curve for extensor digitorum longus (n=3-11). (E) Palmitoyl-L-carnitine (PLC)-induced oxygen consumption in permeabilized fibers (n=3-11). (F) Heatmap of top 100 (50 high and 50 low) genes that were differentially expressed between control and PSD-MKO diaphragms that were reversed in mCATxPSD-MKO diaphragms. Z: z-score. (n=3-4). (G) Heatmap of top 100 (50 high and 50 low) genes that were differentially expressed between control and PSD-MKO diaphragms. Mean ±SEM.

